# Neutrophil-extracellular trap-associated RNA (naRNA) and LL37 enable self-amplifying inflammation in psoriasis

**DOI:** 10.1101/324814

**Authors:** Franziska Herster, Zsofia Bittner, Sabine Dickhöfer, David Eisel, Tatjana Eigenbrod, Thomas Knorpp, Nicole Schneiderhan-Marra, Markus W. Löffler, Hubert Kalbacher, Dominik Hartl, Lukas Freund, Knut Schäkel, Martin Heister, Kamran Ghoreschi, Alexander N.R. Weber

## Abstract

Psoriasis is an inflammatory skin disease with strong immune cell infiltration and high levels of the antimicrobial peptide, LL37. LL37 was shown to complex DNA and RNA to activate plasmacytoid dendritic cells (pDC) in vitro and this has been thought to drive interferon-mediated disease exacerbation. However, the physiological source of such nucleic acids and thus an early initiating event for nucleic-acid mediated inflammation remains unknown. We hypothesized that neutrophils (PMNs), which dominate the psoriatic skin infiltrate, may be the source of DNA and LL37 as well-established components of so-called neutrophil extracellular traps (NETs). We show here that indeed primary murine and human PMNs mount a fulminant NET and cytokine response, albeit not via DNA-but rather RNA-LL37 complexes. This was dependent on TLR8/TLR13 and sensitive to TLR inhibitory oligodeoxynucleotides. Unexpectedly, copious amounts of RNA were identified as a so far unappreciated NET component in vitro and in psoriatic but not healthy skin. Importantly, RNA-LL37-triggered NETs, when transferred to naïve PMNs, prompted additional de novo NET release. Our study highlights TLR-mediated RNA-LL37 sensing by PMNs as parts of a self-propagating vicious cycle that may later engulf pDC and establish chronic inflammation in psoriasis and NET-associated RNA (naRNA) as a physiologically relevant novel component.

**Summary:** Human and bacterial RNA in complex with LL37 activates neutrophils via TLR8 to release cytokines, chemokines and neutrophil extracellular traps (NETs). NETs and neutrophil-rich areas in psoriatic skin contain immunostimulatory RNA, termed here NET-associated RNA (naRNA), and LL37, which may fuel a self-sustaining inflammatory cycle in psoriasis.

## Introduction

Psoriasis is an autoimmune disease of the skin with high incidence in Western countries (1.5-3%), causing high socioeconomic and disease burden with limited but increasing treatment options (Eberle et al., 2016; Griffiths and Barker, 2007). The most common form of psoriasis, plaque psoriasis, is characterized by epidermal hyperplasia due to keratinocyte (KC) hyper-proliferation, increased endothelial proliferation and an infiltrate of leukocytes, such as dendritic cells, T cells and, prominently, polymorphonuclear leukocytes/neutrophils (PMNs) (Griffiths and Barker, 2007). The high and early accumulation of PMNs in psoriatic plaques and micro-abscesses is well documented, as well as an increase of PMNs in the circulation of psoriasis patients, but a specific causative role for PMNs in disease initiation or exacerbation has so far not been defined (Sen et al., 2013) (Schon et al., 2017). Although they are far less abundant in both blood and psoriatic skin, conversely, plasmacytoid dendritic cells (pDCs) have received increased attention in psoriasis. This interest has been fueled by the observations that pDCs can be activated by complexes consisting of either DNA or RNA with a antimicrobial self-peptide that is highly abundant in psoriatic skin, namely LL37: LL37 was shown to form complexes with DNA or RNA that resisted nuclease degradation, were readily taken up by pDCs, and triggered high interferon (IFN) α release from pDCs (Ganguly et al., 2009; Lande et al., 2007). Expression of TLR7 (a pattern recognition receptor for single-stranded RNA) and TLR9 (a receptor for DNA) by pDCs was shown to be vital for the process in vitro.

However, as this scenario of pDC activation strictly requires the *prior* presence of DNA-LL37 and/or RNA-LL37 complexes, the above-described pDC-related mechanism does not qualify as an early initiating event in psoriasis and a process providing the three pDC-stimulating ingredients - LL37, DNA and/or RNA – must be upstream. Unfortunately, the nature of this process has not been discovered to date.

Conversely to pDCs, PMNs can release DNA via so-called neutrophil extracellular trap (NET) formation (NETosis), an activation-induced process leading to the extrusion of nuclear DNA (Brinkmann et al., 2004). In addition, cellular proteins are important components of NETs, and this includes LL37 (Radic and Marion, 2013) for which PMNs are also the main producers in the skin (Sorensen et al., 1997). LL37 is an amphipathic, positively-charged 37 amino acid peptide generated from a precursor protein, the cathelicidin hCAP18 (Griffiths and Barker, 2007; Morizane and Gallo, 2012; Sorensen et al., 2001), that is stored in the secondary granules of PMNs, from where it can be released upon activation (Sorensen et al., 1997).

PMNs thus combine the abilities to release (i) DNA and (ii) LL37 as components of immuno-stimulatory ligand complexes via NETosis, although this has not been firmly linked to psoriasis. They may themselves also sense such ligands via TLRs as expression of TLR8, another RNA sensor (Eigenbrod et al., 2015), and TLR9 (Berger et al., 2012) (Lindau et al., 2013), but not TLR3 or TLR7 (Berger et al., 2012), has been demonstrated but not functionally evaluated. We therefore hypothesized that PMNs may be the source of at least DNA and LL37 as immuno-stimulatory components and that LL37-mediated DNA sensing via TLRs in PMNs might initiate and fuel inflammatory cytokine production and thus inflammation and immune cell infiltration in psoriatic skin.

We here present experimental evidence that human primary PMNs do not sense DNA-LL37 but readily respond to RNA-LL37 complexes, leading to the release of a broad array of chemokines and cytokines and, importantly, NETs, via TLR8 (human) and TLR13 (mouse). Unexpectedly, these NETs contained RNA as a so-far unappreciated component, and they could propagate de novo NETosis in naïve PMNs. PMNs, LL37 and, surprisingly, RNA was also highly abundant in psoriatic but not healthy skin, indicating that PMNs and NET-derived RNA-LL37 complexes may function as integral components of a self-propagating inflammatory cycle.

## Results

### LL37 promotes RNA uptake and induces PMN activation via endosomal TLRs

Previous results indicated that human primary PMNs can respond to RNA and DNA when stimulated for >12 h, albeit at much lower levels than when stimulated with the nucleoside analog TLR7/8 agonist, R848 (Janke et al., 2009; Lindau et al., 2013). We sought to re-evaluate these findings using highly purified primary PMNs assayed within a short time period (4 h) that excludes secondary release effects, e.g. by apoptosis (Fig. S1A). The TLR7/8 agonist R848, like the TLR4 agonist, LPS, elicited robust IL-8 release but only LPS triggered CD62L shedding; phospho-thioate (PTO) synthetic CpG ODN, a typical TLR9 agonist, also strongly activated IL-8 release and CD62L shedding (Fig. 1A, Table S1). However, unmodified, natural phospho-diester DNA ODN or human genomic DNA elicited neither IL-8 release nor CD62L shedding, irrespective of whether they were complexed with LL37 (Fig. 1B). In contrast, in pDCs, LL37 binding of natural DNA triggered potent TLR9 responses (Lande et al., 2007). In the absence of LL37, single-stranded synthetic RNA40 (henceforward referred to as ‘RNA’) barely caused IL-8 release when applied at equimolar concentrations with R848 (Fig. 1C). This suggests that on their own neither RNA nor DNA are able to trigger primary PMN responses. This may be due to the endosomal localization of TLR8 and 9, which R848 and CpG can apparently access, whereas RNA and normal DNA cannot (Kuznik et al., 2011). However, robust IL-8 release and moderate CD62L shedding were observed when RNA was complexed with LL37 (Fig. 1C and cf. Fig. 2E). To check whether PMNs could engage RNA-LL37 complexes, live PMNs were seeded and RNA-LL37 complexes added for 20 min before fixing the cells for electron microscopy analysis. As shown in Fig. 1D, PMNs were found in proximity to fiber-like structures corresponding to RNA-LL37 complexes (Schmidt et al., 2015). Flow cytometry using AlexaFluor (AF) 647-labeled RNA showed that LL37 also promoted the uptake of complexed RNA: >20% of PMNs incubated with labeled RNA in the presence of LL37 were AF647-positive within 60 min, compared to 5% in the absence of LL37 (Fig. 1E). ImageStream bright-field cytometry confirmed that this was not due to external/surface binding of labeled RNA but rather internalization (Fig. 1F, G). Bright-field fluorescence microscopy with Atto488-labeled LL37 showed co-localization of AF647-RNA and Atto488-LL37 in intracellular compartments (Fig. 1H), where endosomal TLRs are expressed (Berger et al., 2012). To pharmacologically explore whether an endosomal TLR might be involved, we investigated the effect of chloroquine, a well-known inhibitor of endosomal TLRs that was non-toxic for PMNs (Fig. S1B). Indeed IL-8 release from primary PMNs in response to RNA-LL37 and CpG ODN (positive control) was significantly reduced by chloroquine addition (Fig. 1I and S1C). As chloroquine does not affect cytoplasmic RNA sensors (e.g. RIG-I) (Matsukura et al., 2007), this implicates an endosomal TLR-dependent in RNA-sensing. We next sought to check whether the observed effect on cytokine release extended to normal RNA from mammalian cells which may be more physiologically relevant in the context of psoriasis. As shown in Fig. 1J, human mRNA significantly induced IL-8 release, but only in combination with LL37. Since microbiome analyses have recently revealed alterations in staphylo- and streptococcal skin colonization in psoriasis patients (Alekseyenko et al., 2013) and bacterial RNA is an emerging immuno-stimulatory pattern (Eigenbrod and Dalpke, 2015), we also tested whether LL37 promoted *S. aureus*, i.e. bacterial RNA (bRNA) activation of PMNs. Fig. 1K shows that bRNA significantly stimulated IL-8 release similarly to RNA40 when complexed with LL37. Taken together this suggests that uptake of RNA is promoted by LL37 and directs RNA to intracellular compartments from which TLR sensing and activation leading to cytokine release can occur – both in response to mammalian and bacterial/foreign RNA.

**Figure 1:**
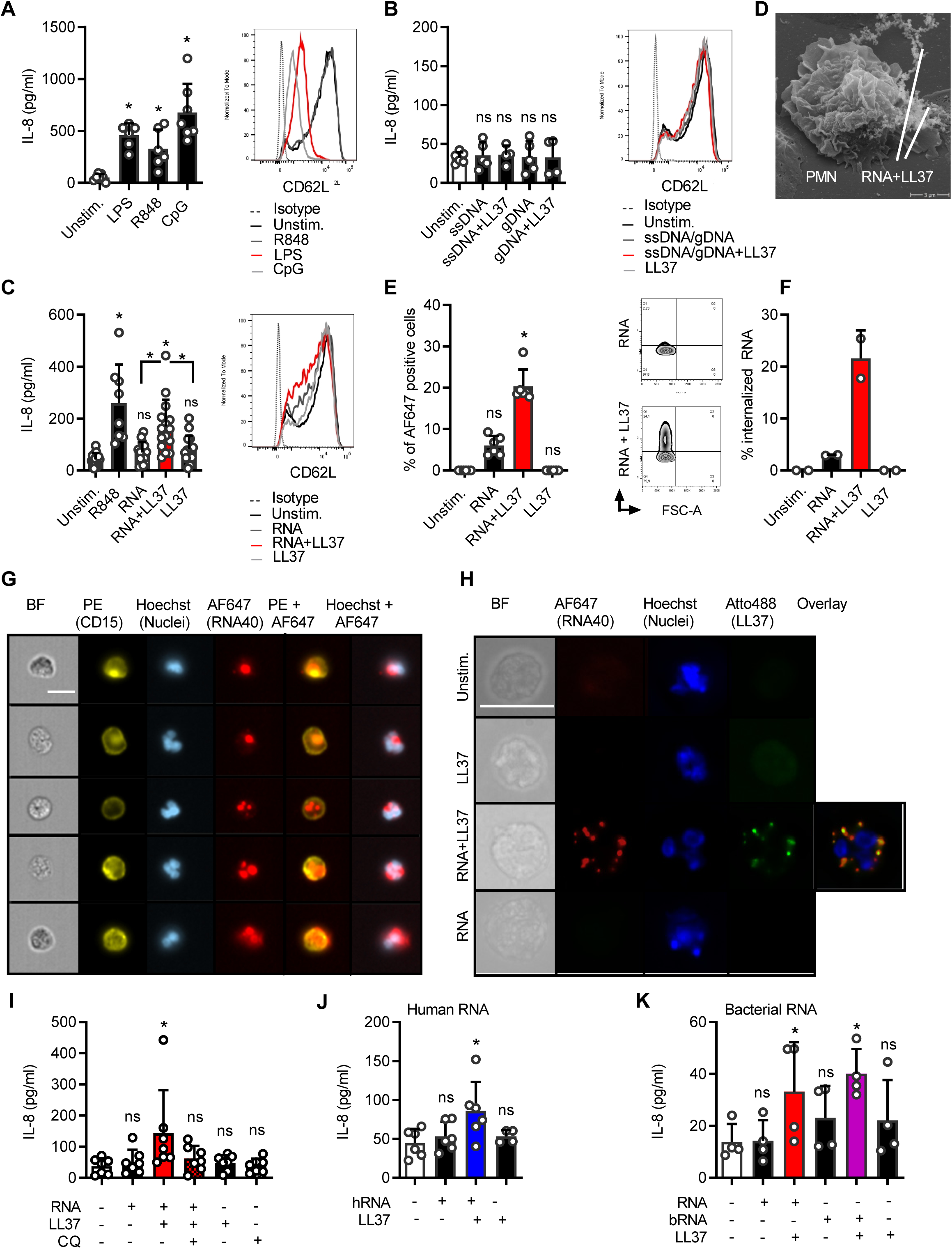
Human neutrophils take up and respond to RNA when complexed to LL37. (A-C) IL-8 release (ELISA, 4 h stimulation) and CD62L shedding (flow cytometry, 1 or 2h) by PMNs from healthy donors stimulated with LPS, R848 or CpG ODN (A, n=6-7), ssDNA and genomic DNA (B, n=5), or RNA40 (C, n=8-15), with or without LL37. (D) Electron microscopy of PMNs incubated with RNA-LL37 for 20 min (n=1). (E-G) FACS (E, n=6), ImageStream cytometry (F, G, scale bar = 10 μm) or conventional bright-field microscopy (H, n=2, scale bar = 10 μm) of PMNs incubated for 60 min with RNA-AF647 complexed with LL37 (n=6). In F the % of cells with ‘internalized’ features are shown (n=2, see Methods), in G selected cells from one representative donor are shown. In H unmodified LL37 was replaced with Atto488-labeled LL37, one representative donor shown. (I) as in C but including chloroquine (CQ) pre-incubation for 30 min (n=6-7). (J) and (K) as in C but using total human RNA from HEK293T cells (J, n=4-6) or bacterial RNA isolated from *S. aureus* (K, n=4) with and without LL37. A-C, E, F, I-K represent combined data (mean+SD) from ‘n’ biological replicates (each dot represents one donor). In D, G and H one representative of ‘n’ biological replicates (donors) is shown (mean+SD). * p<0.05 according to one-way ANOVA with Dunnett’s correction (A, B, C, I, J) or Friedman test with Dunn’s correction (E, K).

**Figure 2:**
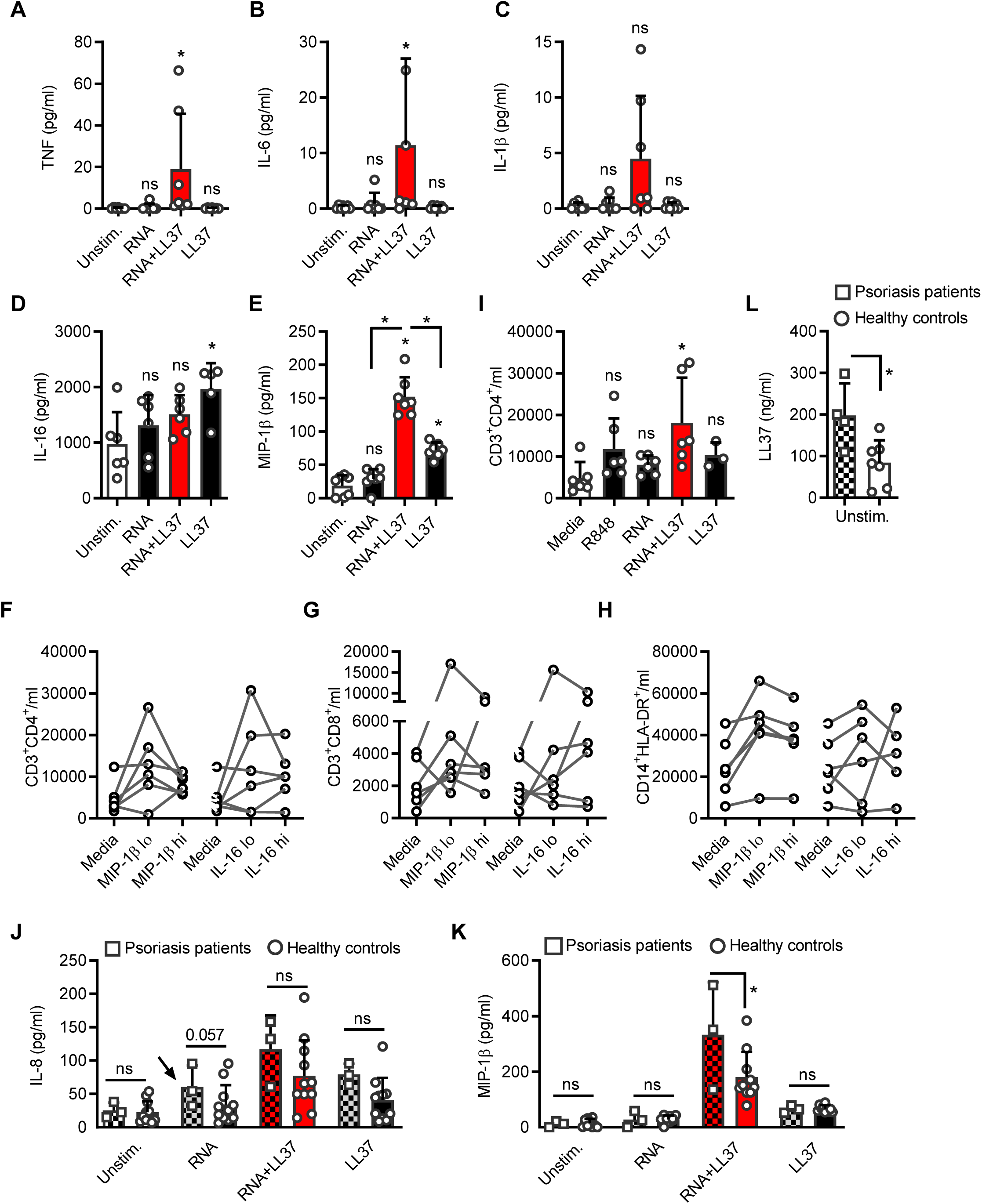
RNA-LL37 complexes promote release of multiple pro-inflammatory cytokines and chemokines, especially in psoriasis PMN. Cytometric bead array (A-C) or ELISA (D, E) for TNF (A), IL-6 (B), IL-1β (C), IL-16 (D) or MIP-1β (E) secreted from PMNs stimulated for 4 h as indicated (n=6). Flow cytometric cell counts of migrated CD4 T cells (F, I), CD8 T cells (G) and CD14^+^HLA-DR^+^ monocytes (H) quantified in transwell assays with total PBMCs in the upper and either MIP-1β (30 and 150 pg/ml) or IL-16 (300 and 1500 pg/ml) (F-H, n=6-7, p>0.05 for treatments vs. media) or R848, RNA with and without LL37 (I, n=3-7) in the lower chamber. (J-K) ELISA of IL-8 and MIP-1β secreted from psoriasis PMNs (n=3 patients, 10 healthy controls, squares, chequered bars) or PMNs from sex-and age-matched healthy donors.(L) as in J or K but ELISA for LL37 (n=4 patients, 7 healthy donors). A-K represent combined data (mean+SD) from ‘n’ biological replicates (each dot represents one donor). * p<0.05 according to Friedmann test with Dunn’s correction (A, B, D-H), or one-way ANOVA with Dunnett’s correction for multiple testing (C, I, J-L).

### RNA-LL37 complexes prompt release of multiple pro-inflammatory cytokines and chemokines, especially in psoriasis PMNs

To test whether RNA-LL37 also promoted the release of cytokines other than IL-8, we screened supernatants of stimulated PMNs by Luminex multiplex analysis and detected TNF, IL-1β, IL-6, and – unexpectedly – IL-16 and MIP-1β primarily or exclusively in RNA-LL37-stimulated cell supernatants, respectively (Fig. S2A,B). Despite considerable variation between donors, this was confirmed by cytometric bead arrays analyses (Fig. 2A-C) and ELISA (Fig. 2D, E) in samples from more donors. To our knowledge, receptor-mediated release of the chemoattractants such as IL-16 (also known as Lymphocyte Chemoattractant Factor, LCF) and MIP-1β (also known as Chemokine (C-C motif) Ligand, CCL4) from primary human PMNs has not been reported. In fact, release of IL-16 was also boosted by LL37 alone (Fig. 2D) and MIP-1β was strongly released from PMNs in response to RNA-LL37 complexes but not RNA alone (Fig. 2E).

Since IL-16 and MIP-1β are known chemoattractants for CD4 T cells and a variety of other immune cells (Cruikshank and Little, 2008; Menten et al., 2002; Roth et al., 2016), we checked whether IL-16 or MIP-1β, at the concentrations observed in this study, could influence the migration and infiltration of additional immune cells, which is a hallmark of psoriatic plaques (Schon et al., 2017). Transwell experiments with peripheral blood mononuclear cells (PBMCs) from healthy human donors (upper well) and IL-16 (300 and 1500 pg/ml), MIP-1β (30 and 150 pg/ml) or SDF-1α (100 ng/ml) as a control (lower well) showed that the lowest concentration of both cytokines induced a donor-dependent increase in the number of CD3+CD4+ helper, CD3+CD8+ cytotoxic T cells as well as CD14+HLA-DR+ monocytes (Figs. 2F-H and S2C-E). Possibly due to a non-linear dose response relationship observed for some cytokines (Atanasova and Whitty, 2012), higher concentrations had a weaker effect in some donors. Owing to the donor-to-donor variation generally observed throughout in the human system, only a non-significant, moderate effect was observed. We separately tested whether RNA or RNA-LL37 in the lower chamber had any direct influence on the migration of these cell populations and noted an unexpected and significant chemoattractive effect on CD4+ T cells, in response to RNA-LL37 (Fig. 2I). Interestingly, the presence of PMNs, regardless of stimulus, was sufficient to attract a variety of other cells (results not shown) due to a so far unknown mechanism.

When repeating the above described stimulation experiments with psoriasis PMNs compared to PMNs from healthy donors, both IL-8 and MIP-1β release was significantly increased two-fold in response to RNA-LL37 (Fig. 2J, K), but not to LPS (Fig. S2F, G). Whereas IL-8 release in response to ‘RNA alone’ (i.e. without LL37) was indistinguishable from ‘unstimulated’ samples in healthy donor PMNs (*cf.* also Fig. 1C), psoriasis PMNs treated with ‘RNA alone’ (Fig. 2J, arrow) produced significantly more IL-8 than both ‘unstimulated’ psoriasis PMNs and healthy donor PMNs stimulated with ‘RNA alone’) Since in healthy donors, RNA seemed to strongly depend on LL37 for efficient PMN uptake and stimulation (*cf*. Fig. 1E, F) and PMNs are the major contributors to elevated LL37 in psoriasis, we speculated whether psoriasis PMNs might constitutively secrete LL37 and this endogenous LL37 might promote an increased responsiveness to exogenously added RNA alone in psoriasis PMNs. Psoriasis PMNs constitutively showed an increased baseline release of LL37 compared to healthy donors (Fig. 2L). We conclude that the combination of RNA with LL37 triggers the release of an extended array of pro-inflammatory cytokines and chemoattractants not triggered by RNA alone. This was more prominent in psoriasis PMNs, potentially due to a higher abundance of PMN-derived LL37.

### RNA-LL37 complexes trigger the release of RNA- and LL37-containing NETs (naRNA)

Our data so far indicate that RNA and LL37 might contribute to cytokine-mediated inflammation and immune infiltration in neutrophil-containing skin lesions in psoriasis patients. Such neutrophil responses would probably be temporary, unless RNA-LL37 triggered the release of additional RNA and LL37. PMN NETs are known to contain extracellular DNA and DNA can act as an immune stimulant of pDCs when complexed with LL37 (Lande et al., 2011). But DNA-LL37 cannot activate PMNs (*cf.* Fig. 1B) and whether RNA is released by NETosis has not been shown. NETs would thus only propagate PMN activation in case they also contained RNA. We therefore next tested (i) whether RNA-LL37 induces NETosis and (ii) whether NETs contain RNA. Interestingly, electron microscopy (Fig. 3A) and a elastase-based assay (see Methods) confirmed significant neutrophil elastase release, a hallmark of NETosis, in response to RNA-LL37 and PMA (positive control), but also with LL37 alone (Fig. 3B). However, fluorescence microscopy analysis of fixed PMN samples established that only RNA-LL37 complexes, and not LL37 alone, prompted the formation of LL37-positive NET-like structures (Fig. 3C). To check whether the NETs also contained RNA, an RNA-selective fluorescent dye, SYTO RNAselect, was used to stain NETs. This staining showed released endogenous RNA (green) in both PMA and RNA-LL37-mediated NETs (Fig. 3C). Although the dye’s specificity for RNA has been confirmed already (Li et al., 2006) we confirmed that RNA staining, but not DNA staining, was sensitive to RNase A and thus RNA-specific (Fig. S3A, B). The presence of cellular RNA in NET structures was also confirmed by the use of specific Abs against pseudo-uridine (ΨU), a nucleotide absent from both DNA and also the synthetic RNA40 used for LL37 complex formation and stimulation (Fig. 3D). Confirming the specificity of the stain, the signal was completely RNase sensitive. Furthermore, using AF647- or AF488-labeled synthetic RNA for in vitro stimulation both the extracellular SYTO RNAselect and anti-ΨU signals could be unequivocally attributed to de-novo released ‘cellular RNA’ and distinguished from exogenously added ‘stimulant (synthetic) RNA’ by (Fig. S3C, D). The same applied to LL37, using Atto488-LL37 (Fig. S3E). To further exclude artefacts that might arise from staining fixed samples, RNA-LL37 mediated NETosis was also analyzed and confirmed using live-cell time-lapse analysis of PMNs (Fig. 3E and Movies S1-S3, quantified in Fig. 3F). Interestingly, in PMNs captured on the verge of NETosis, RNA staining in a granula-like fashion could be clearly observed (Fig. S3F). Since increased levels of NETs containing DNA have already been reported in the blood and skin of psoriasis patients (Hu et al., 2016), we next investigated whether this was also the case for RNA. We therefore stained skin sections with SYTO RNAselect, anti-LL37 and anti-NE Abs. Indeed, psoriatic skin was highly positive for LL37; interestingly, the RNA signal was also generally stronger when compared to healthy skin, using identical staining and microscopy settings (Fig. 3G). RNA and LL37 were frequently co-localized in samples with high PMN infiltration (anti-NE staining), (Fig. 3H and Movie S4). SYTO RNAselect specificity was also confirmed in skin sections using anti-ΨU staining with both stains clearly overlapping (Fig. S3G). This indicates that RNA-LL37 complexes may act as physiologically relevant activators in psoriatic skin. Further, the ability of RNA-LL37 to drive primary PMNs to extrude NETs containing RNA in addition to DNA and LL37 may represent the so far unknown activating event upstream of pDC activation.

**Figure 3:**
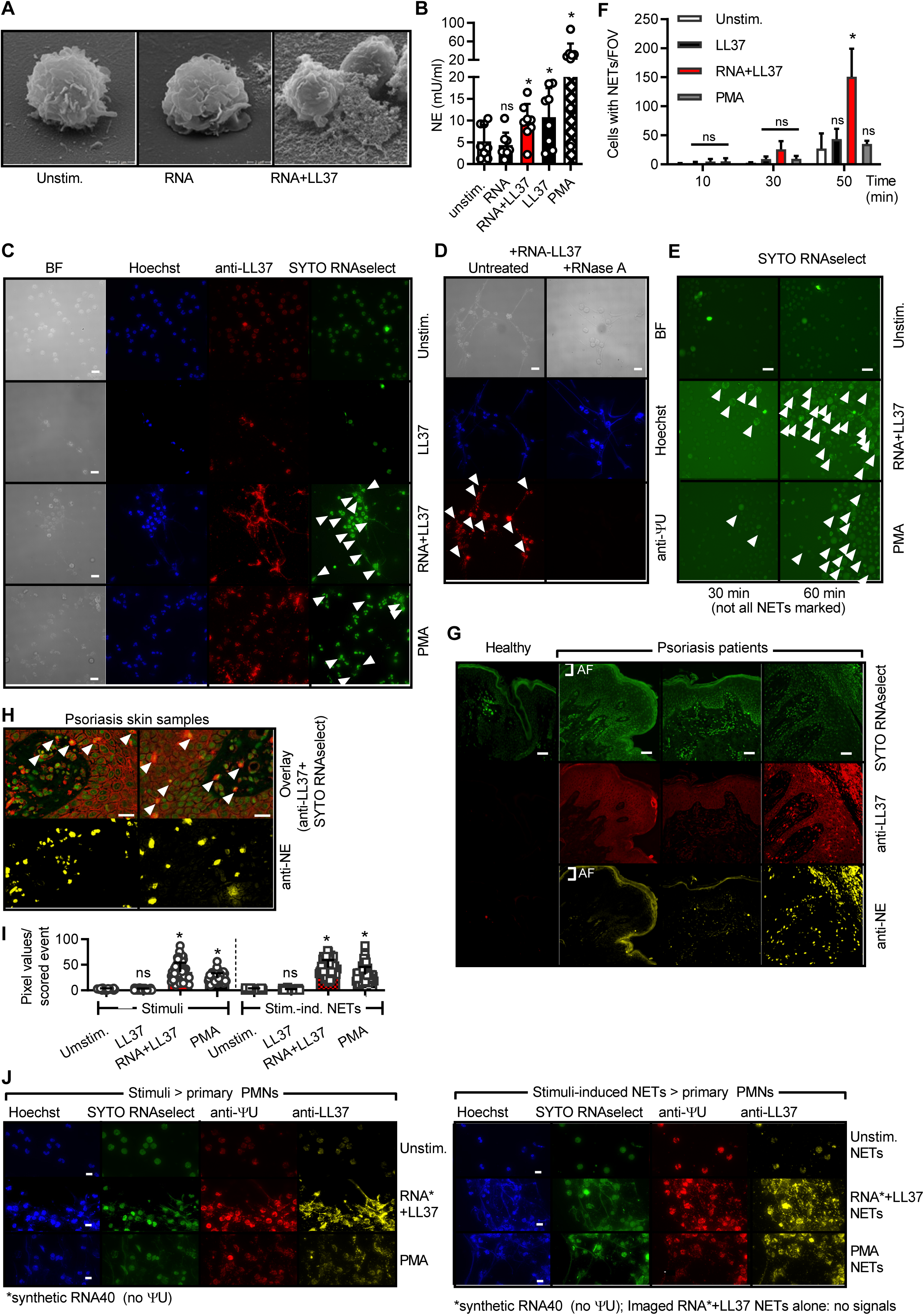
RNA-LL37 trigger the release of NETs containing further RNA, DNA and LL37. (A) EM pictures from PMNs stimulated with RNA-LL37. (B) Neutrophil elastase (NE) release from PMNs stimulated for 3 h (n=8, each dot represents one donor). Fluorescence microscopy of fixed and Hoechst, anti-LL37, SYTO RNAselect- (C) and/or anti-ΨU-stained (D, with or without RNase A treatment) or live (E) PMNs stimulated as indicated (n=6, scale bar = 10 µm or n=4, scale bar = 20 µm). (F) Quantification of live cell microscopy), see also Movies S1-S3. (G, H) Skin sections from healthy (G) or psoriasis- (G, H) affected skin (n=12 patients and 3 healthy controls, scale bar = 20 µm). (I, J) as in C (left panels) and stimulated with transferred NETs (right panels). AF = autofluorescence. B, F and I represent combined data (mean+SD) from ‘n’ biological replicates. In A, C-H and J representative samples of ‘n’ replicates or donors are shown. Arrowheads indicate released RNA/NETs (C-E) or RNA-LL37 co-localization (H). * p<0.05 according to one-way ANOVA (B), two-way ANOVA (F) with Dunnett’s correction or Kruskal-Wallis test with Dunn’s correction (I).

Based on these results it also appeared plausible that RNA-LL37 complexes might trigger a NET-mediated self-propagating inflammatory loop of repetitive PMN activation, in which NET material would act in a manner similar to RNA-LL37 and activate unstimulated/naïve PMNs to undergo NETosis, leading to repetitive cycles of immune activation. Indeed, when we transferred NETs harvested from PMNs stimulated with PMA or RNA-LL37 complexes to unstimulated/naïve PMNs, these responded with de-novo NET release in turn (Fig. 3J, quantified in I). Conversely, material harvested from LL37 or untreated PMNs, which did not contain NETs, did not. Collectively, NETs generated by initial RNA-LL37 activation had the capacity to activate naive PMNs to extrude further DNA, RNA and LL37, thus providing the requirements for a self-propagating feed forward loop of immune activation that might apply to skin PMNs. At the same time, we provide evidence for the so far unappreciated existence and physiological relevance of RNA in NETs, which we would like to term NET-associated RNA or naRNA.

### RNA-LL37 complexes and NETs activate PMNs via TLR8/TLR13

In order to gain an insight which receptor mediates the observed effects of RNA-LL37 in primary PMNs, we first compared BM-PMNs from WT mice and mice deficient for *Unc93b1*, a critical chaperone of endosomal TLRs (Eigenbrod et al., 2012). Evidently, not only the cytokine response of BM-PMNs to bRNA-LL37 (and the control stimulus CpG) was strictly dependent on *Unc93b1* and thus endosomal TLRs (Fig. 4A); NETosis triggered by bRNA-LL37 complexes, but not the TLR-independent control stimulus PMA, was also *Unc93b1* dependent (Fig. 4B).Given that RNA is a known dual TLR7 and TLR8 agonist (Colak et al., 2014) and primary human PMNs do not show *TLR7* expression (Berger et al., 2012), we suspected TLR8 to mediate RNA-LL37-mediated PMN activation. Unfortunately, human primary PMNs are not amenable to RNAi or genetic editing. Therefore, we first tested the responsiveness of BM-PMNs from mice deficient for TLR13, the functional counterpart of human TLR8 and primary bRNA sensor in mice (Eigenbrod and Dalpke, 2015). In murine PMNs both cytokine and NET responses to bRNA-LL37 were strictly dependent on TLR13 (Fig. 4A, C), whereas the negative controls LPS, *E. c*oli or PMA were not, respectively. Complementarily, the failure of *TLR8* CRISPR-deleted BlaER1 monocytes (Vierbuchen et al., 2017) to respond to RNA-LL37 complexes (Fig. 4D) indicated a dependence of RNA-LL37 on TLR8 in the human system. Thus the stimulatory potential of RNA-LL37 complexes can be considered to be dependent on the TLR8/TLR13 RNA sensing system (Eigenbrod et al., 2015).

**Figure 4:**
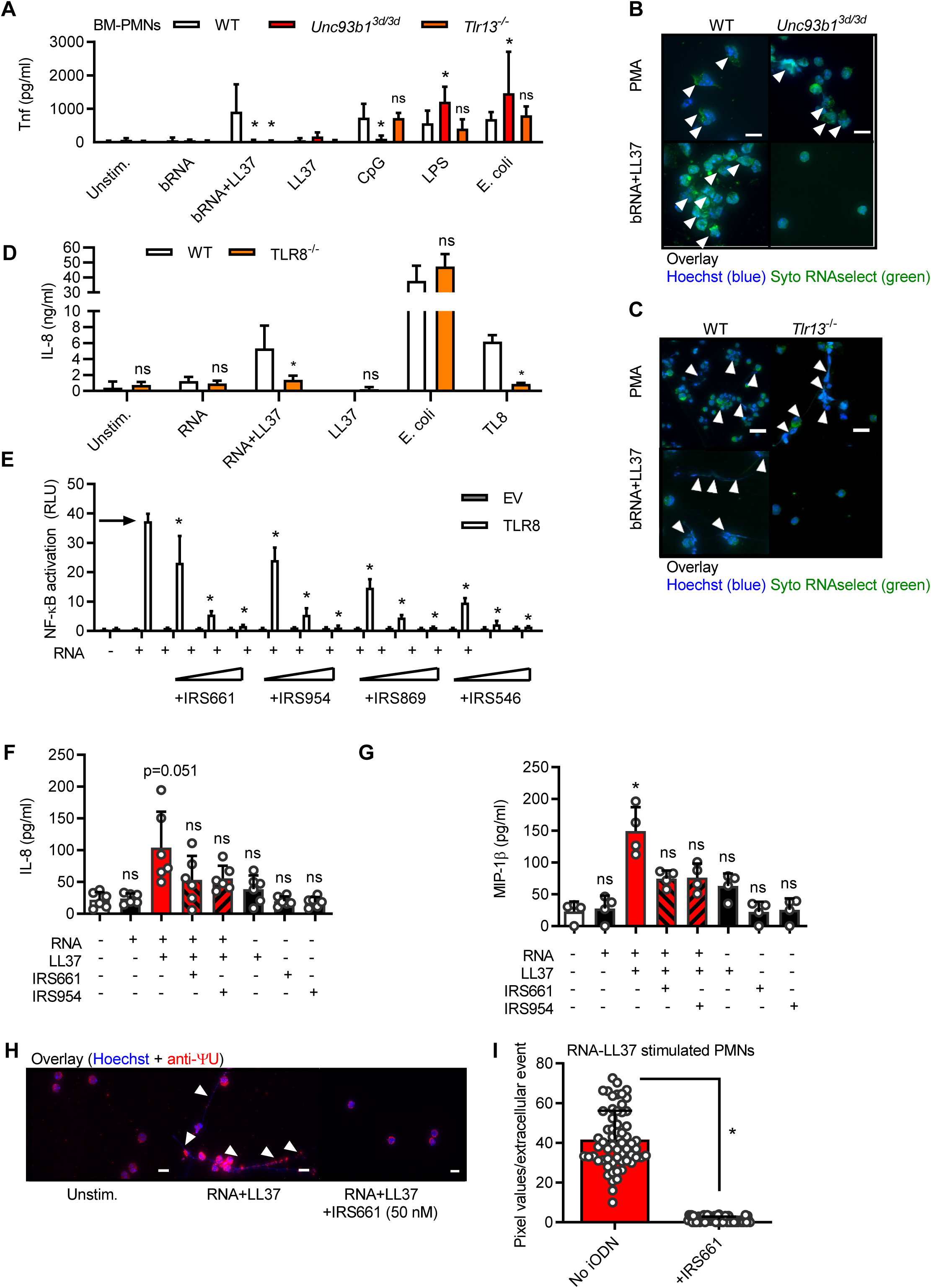
RNA-LL37 effects in primary PMNs are mediated by TLR8/TLR13 and can be inhibited by Inhibitory oligodeoxynucleotides. (A) Tnf release (ELISA, 5 h stimulation as indicated) by BM-PMNs from different mouse strains (n=4-8 each). (B, C) Fluorescence microscopy of fixed and Hoechst and SYTO RNAselect-stained stimulated BM-PMNs from different mouse strains (n=4 Tlr13^-/-^, n=5 Unc93b1^3d/3d^, n=8 WT; scale bar = 10 µm or n=4, scale bar = 20 µm). (D) IL-8 and TNF release (ELISA, 18 h stimulation as indicated) by WT and *TLR8* CRISPR-edited BlaER1 cells (n=7). (E) NF-κB dual luciferase reporter assay in HEK293T cells, transfected with NF-κB firefly luciferase reporter, *Renilla* control reporter and TLR8 plasmid and subsequently stimulated with RNA without (arrow) or with IRS661, IRS954, IRS869 and IRS546 (0.15-3.5 µM, n=2 each). IL-8 (F, n=6) or MIP-1β (G, n=4) release from stimulated PMNs with or without IRS661 (1 nM) or IRS954 (50 nM) pre-incubation (30 min) quantified by ELISA. (H) Fluorescence microscopy of fixed and Hoechst and anti-ΨU-stained RNA-LL37-stimulated PMNs (n=3, scale bar = 10 µm). (I) Quantification of H. A, D, E, F, G and I represent combined data (mean+SD) from ‘n’ biological replicates (each dot represents one mouse or donor). In B, C, E and H one representative of ‘n’ replicates is shown (mean+SD of technical triplicates). * p<0.05 according to two-way ANOVA with Dunnett correction for multiple testing (E, comparison against ‘no inhibitor’ condition, arrow), one-way ANOVA with Sidak correction (A, F) or Friedmann test with Dunn correction (G).

### PMN cytokine release and NET formation by RNA-LL37 can be blocked by iODNs

Having identified TLR8 as the responsible receptor in humans, we next wondered whether PMN activation by RNA-LL37 could be blocked at the level of TLR activation by so-called inhibitory oligonucleotides (iODNs). IRS869, IRS661 and IRS954 represent such TLR7 and 9-inhibitory iODNs that were recently proposed for the treatment of another autoimmune disease, namely systemic lupus erythematosus (SLE), whereas IRS546 was used as a non-inhibitory ‘control’ ODN (Barrat and Coffman, 2008; Barrat et al., 2005; Duramad et al., 2005; Guiducci et al., 2010). Given the similarities between endosomal TLRs (Colak et al., 2014), we speculated that some of these iODNs could potentially also block TLR8, but the effects of iODNs on this receptor have not been studied systematically. HEK293T cells do not express TLR7-9 but can be rendered responsive to their agonists by exogenously expressing TLR7, TLR8 or TLR9 upon transfection (Colak et al., 2014). In HEK293T cells transfected and stimulated in this way, IRS869 and IRS954 blocked NF-κB activation in response to R848 for TLR7 and 8 (Figs. S4A and B) and CpG for TLR9 (Fig. S4C). IRS661 effectively blocked TLR7-but not TLR9-mediated activation. IRS661 strongly increased R848-mediated NF-κB activation via TLR8 (Fig. S4B), similar to observations made for T-rich ODNs combined with R848 (Jurk et al., 2006) and highlighting the differences between R848 and RNA reported previously (Colak et al., 2014). However, for RNA complexed with DOTAP (a synthetic complexing agent promoting endosomal uptake (Yasuda et al., 2005), similarly to LL37), all three iODNs blocked TLR8-mediated NF-κB activation (Fig. 4E). As the ‘control’ ODN IRS546 blocked NF-κB activation in response to TLR7-9 stimulation, in our hands it did not represent a proper control unlike previously published (Barrat et al., 2005). Nevertheless, the effects of all iODNs seemed specific for endosomal TLR signaling as the effect of all iODNs on NF-κB activity induced by TNF (a non-TLR receptor pathway activating NF-κB, Fig. S4D) or MyD88 overexpression (TLR receptor-independent NF-κB activation, Fig. S4E) were minor. Having excluded a toxic effect on primary PMNs (Fig. S4F), we investigated whether the observed effects could be transferred to endogenous TLR8 activation in primary PMNs, where LL37 as a physiologically relevant uptake reagent was again used. Here, we found that iODNs IRS661 and IRS954 were able to inhibit IL-8 release by primary PMNs from healthy donors in response to RNA-LL37 at nanomolar iODN concentrations so that IL-8 levels were not statistically above background (Fig. 4F). On the other hand, LPS-mediated IL-8 release was unaffected (Fig. S4G and (Saitoh et al., 2012)). The blocking effect extended to MIP-1β (Fig. 4G), similar to the effect observed for chloroquine (Fig. S4H). Importantly, NETosis by primary human PMNs could be effectively blocked by nanomolar concentrations of IRS661 (Fig. 4H, quantified in Fig. 4I). These results are in agreement with the observed TLR8-dependence and iODN-mediated inhibition of NETosis in HIV infections (Saitoh et al., 2012) and indicate an applicability to RNA-LL37 complexes. In conclusion, known pre-clinical iODNs are able to block RNA-LL37-mediated cytokine and NET release.

## Discussion

Although systemic therapies with biologicals have brought benefit for many psoriasis patients in industrialized countries (Eberle et al., 2016), psoriasis remains a significant health problem worldwide. One reason is that early triggers and mechanisms leading to chronic disease, exacerbation and flare-ups are not well understood. Effects of DNA-/RNA-LL37 complexes on pDCs were reported as a novel mechanism in psoriasis inflammation (Ganguly et al., 2009; Lande et al., 2007). However, since pDCs are only a minor cell population even in psoriatic skin and they cannot release nucleic acids or LL37, their contribution to cell-mediated inflammation and the provision of self-ligands had to be restricted to amplifying an already ongoing pathology, characterized by pDC-attracting chemokines and an existing presence of self-ligands in the skin. This begged the question of how nucleic acid-LL37 complexes may be released in the skin to trigger subsequent immune activation. Since PMNs can release nucleic acids, constitute the main sources of LL37 in humans and congregate in psoriatic skin, we chose to study PMN RNA-LL37-mediated responses after TLR8 was identified as an RNA receptor on PMNs (Berger et al., 2012). We now provide novel evidence that PMNs are a likely early source of DNA, RNA and LL37 that may be at the heart of inflammation in situ.

We here found that the combination of RNA and LL37 produces an inflammatory ligand that is not only sensed by PMNs but it also prompted the release of further components for additional RNA-LL37 and DNA-LL37 complexes from PMNs. Whereas for pDCs both “inflammatory ingredients”, RNA or DNA and LL37, required the presence of another cellular source first, a small number of activated and NETing PMNs – by releasing both, LL37 and RNA/DNA – may spark potent, self-propagating inflammation amongst PMNs and *in situ.* Following an initial activation of PMNs by endogenous RNA from damaged skin cells (so-called Köbner phenomenon), immune activation may get out of control due to chemokine release and NET formation (and thus nucleic acid and LL37 extrusion (Neumann et al., 2014a) which has been observed in the skin of psoriasis patients (Hu et al., 2016). Although this will require further investigation, our data for the first time raise the possibility that not only self-RNA, as previously proposed (Ganguly et al., 2009), but also foreign, e.g. bacterial RNA (Eigenbrod and Dalpke, 2015) (*cf.* Fig. 1K) or possibly fungal RNA may unfold immunostimulatory properties in the presence of LL37 triggered during minor skin injury. Although the concentration of the cytokines released after 4 h stimulation was moderate, release over longer time periods and the sheer number of PMNs found in psoriatic skin may nevertheless contribute substantially to local inflammation and attraction of additional leukocytes over time. The PMN-mediated ongoing release of NET DNA, RNA and LL37 would eventually enable pDCs and other immune cells to join the vicious cycle of activation fueled by endogenous nucleic acids (Chamilos et al., 2012; Ganguly et al., 2009; Lande et al., 2007). Although KC- and pDC-released type I IFNs contribute to psoriasis (Afshar et al., 2013; Yao et al., 2008), TNF and IL-1β are additionally required for immune cell infiltration and T cell polarization, respectively (Ghoreschi et al., 2007) and RNA-LL37-mediated activation of PMNs may be the so far missing source of these cytokines. Subsequent cytokine-mediated inflammation is known to favor hyper-proliferation (Hanel et al., 2013), LL37 production (Schauber et al., 2007), TLR9 responsiveness (Morizane et al., 2012) and IFN production (Zhang et al., 2016) in KC, as well as T cell and monocyte attraction and polarization (Ghoreschi et al., 2007). Our study thus raises the possibility that self-amplifying inflammation mediated by RNA and LL37 via TLRs recognition and NETosis of PMNs may represent an early and vital step in psoriasis development (Fig. S5).

An unexpected and novel finding that appears vital for this process and may have wide ramifications in many other NET-related processes, is the observation that RNA is an abundant component of mammalian NETs. Several hundred reports on NETs have so far not provided experimental evidence that NET-associated RNA (naRNA) of physiological importance, let alone is contained in NETs. Our data now demonstrate that naRNA is an abundant component of both RNA-LL37 and PMA-triggered NETs. The observation that it also triggers further immune activation of PMNs additionally demonstrates physiological relevance. NaRNA could thus represent as a novel immunostimulatory component within NETs, which has not been appreciated so far. Interestingly, insect NETs contain primarily RNA rather than DNA (Altincicek et al., 2008), and eosinophils were reported to pre-store RNA within their granules (Behzad et al., 2010), possibly for later extrusion. Of course, free RNA is not stable over long periods, but LL37 may confer protection not only for NETs in general (Neumann et al., 2014b) but also naRNA. NaRNA may thus be a common and functionally important NET component which remains to be explored further. Given that extracellular RNA can also amplify responses to other PRR ligands (Noll et al., 2017), the role of naRNA will thus be interesting to study in other NET-related diseases, e.g. SLE (Gestermann et al., 2018), atherosclerosis (Warnatsch et al., 2015) and even cancer (Gregoire et al., 2015), where the focus so far has been exclusively on NET DNA. Furthermore, given that NETosis first and foremost has been described as an important host-defense mechanism (Brinkmann et al., 2004), it remains to be investigated whether naRNA also executes or participates in antimicrobial activities.

In conclusion, our study offers insights into how PMN RNA-LL37 sensing and NET propagation may contribute to early disease development and, based on the observed effects of iODNs on PMN activation, warrant the further exploration of TLR pathways and PMNs as targets for restricting innate immune activation in psoriatic skin. Additionally, our study also calls for an exploration of naRNA as a novel and intriguing new component of NETs in psoriasis and other processes.

## Materials and Methods

### Reagents

All chemicals were from Sigma unless otherwise stated. PRR agonists and LL37 were from Invivogen except RNA40 (iba Lifescience, normal and AF647- or AF488-labeled) and CpG PTO 2006 (TIB Molbiol), see Table S2 and 3. iODNs used in this study with their respective sequences are listed in Table S3 and were from TIB Molbiol. Total human mRNA was isolated from HEK293T cells using the RNeasy kit on a QIAcube, both from QIAGEN. *S. aureus* RNA was isolated as described (Eigenbrod et al., 2015). Genomic human DNA was isolated from whole blood using QIAamp DNA Blood Mini Kit from Qiagen (51106) and phosphodiester DNA from TIB Molbiol. LL37 was from InvivoGen (see Table S2) and DOTAP from Roth, L787.2. LL37 was Atto488 (from Atto-Tec as a carboxy-reactive reagent)-labeled using standard procedures. For complex formation 5.8 µM RNA40 (approximately 34.4 µg/ml and equimolar to R848 used in this setting), 1 µM ssDNA (approximately 20 µg/ml and equimolar to CpG used in this setting, sequence see Table S3), genomic DNA (20 µg/ml) or bacterial RNA (10 µg/ml) was mixed together with 10 µg LL37 (see Table S2, Atto-488 where indicated) and left for one hour at RT. For experiments with BM-PMNs, 1.25 µg bacterial RNA was complexed with 2.5 µg LL37. For the RNA-only or LL37-only conditions, the same amounts and volumes were used replacing one of the constituents by sterile, endotoxin-free H2O. Ficoll was from Millipore. Antibodies used for flow cytometry, ImageStream analysis or fluorescence microscopy are listed in Table S4 as well as the recombinant cytokines used in this study. Constructs used for HEK293T transfection are listed in Table S5.

### Study participants and sample acquisition

All patients and healthy blood donors included in this study provided their written informed consent before study participation. Approval for use of their biomaterials was obtained by the local ethics committees at the University Hospitals of Tübingen and Heidelberg, in accordance with the principles laid down in the Declaration of Helsinki as well as applicable laws and regulations. All blood or skin samples obtained from psoriasis patients (median age 41.8 years, PASI >10, no systemic treatments at the time of blood/skin sampling) were obtained at the University Hospitals Tübingen or Heidelberg, Departments of Dermatology, respectively, and were processed simultaneously with samples from at least one healthy donor matched for age and sex (recruited at the University of Tübingen, Department of Immunology). Skin sections were from 12 patients with Plaque Psoriasis and 1 patient with Psoriasis guttata.

### Mice and isolation of bone-marrow derived PMNs (BM-PMNs)

*Unc93b1*^3d/3d^-(Tabeta et al., 2006), *Tlr13*-deficient (Li and Chen, 2012) mice (both C57BL/6 background) and WT C57BL/6 mice between 8 and 20 weeks of age were used in accordance with local institutional guidelines on animal experiments and under specific locally approved protocols for sacrificing and *in vivo* work. All mouse colonies were maintained in line with local regulatory guidelines and hygiene monitoring. Bone-marrow (BM)-PMNs were isolated from the bone marrow using magnetic separation (mouse Neutrophil isolation kit, Miltenyi Biotec, 130-097-658) and following the manufacturer’s instructions. PMN stimulation was carried out for 5 hours at 37 °C and 5% CO_2_. Thereafter supernatants were harvested and used for ELISA. For microscopy the cells were stimulated for 16 h and subsequently stained.

### Neutrophil isolation and stimulation

Whole blood (EDTA-anticoagulated) was diluted in PBS (Thermo Fisher, 14190-169), loaded on Ficoll (1.077 g/ml, Biocoll, ab211650) and centrifuged for 25 min at 506 x g at 21 °C without brake. All layers were discarded after density gradient separation except for the erythrocyte-granulocyte pellet. Thereafter, erythrocyte lysis (using 1x ammonium chloride erythrocyte lysis buffer, see Table S6) was performed twice (for 20 and 10 min) at 4 °C on a roller shaker. The remaining cell pellet was carefully resuspended in culture medium (RPMI culture medium (Sigma Aldrich, R8758) + 10% FBS (heat inactivated, sterile filtered, TH Geyer, 11682258)) and 1.6 x 10^6^ cells/ml were seeded (24 well plate, 96 well plate). After resting for 30 min at 37°C, 5% CO_2_, the cells were pre-treated with inhibitors (where indicated) for 30 min and subsequently stimulated with the indicated agonists for 4 hours (for ELISA) or for 30 min to 3 h (for FACS analysis or microscopy).

### PBMC isolation

Whole blood (EDTA anticoagulant) was diluted in PBS. After density gradient separation using Ficoll (described above), the PBMC layer was then carefully transferred into a new reaction tube and diluted in PBS (1:1). The cell suspension was spun down at 645 x g for 8 min. The cells were then washed twice more in PBS, resuspended in culture medium (RPMI + 10% FBS (heat inactivated) + 1% L-glutamine), before counting and seeding.

### BlaER1 cells culture, transdifferentiation and stimulation

BlaER1 cells (WT and TLR8 -/-, kindly provided by Tatjana Eigenbrod from Heidelberg) were cultured in culture medium (VLE-RPMI (Merck Biochrom, FG1415) + 10 % FBS, 1 % Pen/Strep (Gibco, 15140122), 1 % sodium pyruvate (100 mM, Gibco, 11360070), 1 % HEPES soluation (Sigma, H0887)). After reaching a cell concentration not higher than 2 x 106/ml the cell were seeded in a 6 well plate (1 x 106 cells per well) and transdifferentiated in culture medium adding 150 nM β-estradiol, 10 ng/ml M-CSF and 10 ng/ml hIL-3 for 6 days including 2 medium changes (on day 2 and 5). On day 7 the adherent cells were counted again, seeded (96 well plate, 5 x 104 cells per well in culture medium) and left to rest for 1h. Subsequently, they were stimulated for 18 h and the supernatants were harvested and collected for ELISA measurements. The transdifferentiation efficiency was verified by FACS analysis, using CD19, CD14 and CD11b as cell surface markers.

### Flow cytometry

After PMN isolation and stimulation, the purity and activation status of neutrophils was determined by flow cytometry. 200 µl of the cell suspension was transferred into a 96 well plate (U-shape) and spun down for 5 min at 448 x g, 4 °C. FcR block was performed using pooled human serum diluted 1:10 in FACS buffer (PBS, 1 mM EDTA, 2% FBS heat inactivated) for 15 min at 4 °C. After washing, the samples were stained for approximately 20-30 min at 4°C in the dark. Thereafter, fixation buffer (4% PFA in PBS) was added to the cell pellets for 10 min at RT in the dark. After an additional washing step, the cell pellets were resuspended in 150 µl FACS buffer. Measurements were performed on a FACS Canto II from BD Bioscience, Diva software. Analysis was performed using FlowJo V10 analysis software.

### ELISA

Cytokines were determined in half-area plates (Greiner, Bio-one) using duplicates or triplicates and measuring with a standard plate reader. The assays were performed according to the manufacturer’s instructions (Biolegend, R&D systems), using appropriate dilutions of the supernatants. For LL37 determination a kit from HycultBiotech (HK321-02) was used following the manufacturer’s instructions.

### ImageStream analysis

ImageStream analysis was used to analyze internalization of RNA-LL37 complexes using spot-counts and tracking single cells. The cells were first seeded in a 96 well plate, 8 x 10^6^ cells/ml, 125 µl per well. Subsequently, they were stimulated for 1 hour with RNA-AF647 (IBA technologies) and/or LL37-Atto488 (kindly provided by Hubert Kalbacher, University of Tübingen). FcR block and surface staining (here CD15 PE) was performed as described above. After fixation, the cells were permeabilized with 0.05 % Saponin (Applichem, A4518.0100) for 15 min at RT in the dark. After washing, nuclei were stained with Hoechst 33342 (Sigma, B2261, 1 µg/ml) for 5 min at RT in the dark, washed and resuspended in 50 µl FACS buffer and transferred into a 1.5 ml Eppendorf tube. At least 10.000 cells were acquired for each sample with 40x magnification using an ImageStream X MKII with the INSPIRE instrument controller software (Merck-Millipore/Amnis). Data were analyzed with IDEAS Image analysis software. All samples were gated on single cells in focus.

### Fluorescence Microscopy of fixed neutrophils

The cells were seeded in a 96 well plate at 1.6 x 10^6^ cells/ml, 125 µl per well. Subsequently they were stimulated with the complexes for 30 min and 1 h using RNA-AF647/AF488 and/or LL37-Atto488. FcR block, staining, fixation and permeabilization were performed as for Flow cytometry. The cell pellets were resuspended in 50-100 µl FACS buffer. 40 µl of the cell suspension was pipetted on a Poly-L-Lysine coated coverslip (Corning, 734-1005) and the cells were left to attach for one hour in the dark. ProLong Diamond Antifade (Thermo Fisher, P36965) was used to mount the coverslips on uncoated microscopy slides. For NET analysis PMNs were seeded in 24 well plates, containing Poly-L-Lysine coated coverslips and stimulated with RNA-LL37 complexes or PMA (600 nM) for 3 hours. NETs were fixed and stained using the protocol from Brinkmann et al.(Brinkmann et al., 2010). Where indicated, 100 µg/ml RNase A (DNase, protease-free, ThermoFisher, EN0531) was added after fixation and incubated overnight at 37°C. RNA was stained using SYTO RNAselect Green fluorescent dye (Thermo Fisher, 50 µM) or anti-ΨU antibody (see Table S4) and nuclear DNA was stained with Hoechst33342 (Thermo Fisher, 1 µg/ml). LL37 and PMNs were visualized using an unconjugated rabbit anti-LL37 antibody or unconjugated mouse anti-NE with subsequent staining with an AF647-conjugated anti-rabbit or an AF594-conjugated anti-mouse antibody, respectively (see Supplementary Table S4), after blocking with pooled human serum (1:10 in PBS). Secondary antibodies alone did not yield any significant staining. The slides were left to dry overnight at RT in the dark and were then stored at 4 °C before microscopy. The measurements were conducted with a Nikon Ti2 eclipse (100x magnification) and the analysis was performed using Fiji analysis software.

### Fluorescence Microscopy of tissue samples

Skin samples from psoriasis patients with a PASI ≥ 10 and without systemic treatment, and healthy skin samples paraffin-embedded according to standard procedures were deparaffinized and rehydrated using Roti Histol (Roth, 6640.1) and decreasing concentrations of ethanol (100%, 95%, 80% and 70%). After rinsing in ddH_2_O, antigen retrieval was performed by boiling for 10-20 min in citrate buffer (0.1 M, pH=6). The skin tissue was then washed 3 times for 5 min with PBS. Blocking was performed using pooled human serum (1:10 in PBS) for 30 min at RT. The primary antibody was added either overnight at 4°C or for 1 hour at RT. After 3 washes, the secondary antibody was added for 30 min at RT in the dark. After another 3 washes, SYTO RNAselect Green fluorescent dye (Thermo Fisher, 50 µM) was added for 20 min at RT in the dark. Thereafter, the samples were washed again and Hoechst 33342 (ThermoFisher, 1 µg/ml) was added for 5 min. Then 3 last washes were performed before using ProLong Diamond Antifade (Thermo Fisher, P36965) for mounting. The samples were left to dry overnight at RT in the dark before being used for microscopy or stored at 4°C. The specimens were analyzed on a Nikon Ti2 eclipse bright-field fluorescence microscope (10x-60x magnification) and the analysis was performed using Fiji analysis software. Autofluorescence in multiple channels typical for the stratum corneum was labeled “AF”.

### Live cell imaging of primary neutrophils

Human neutrophils were isolated by magnetic separation using MACSxpress whole blood neutrophil isolation kit (Miltenyi Biotec, 130-104-434) and were seeded into a micro-insert 4 well dish (Ibidi 80406). Hoechst 33342 (1 µg/ml) and SYTO RNAselect Green fluorescent dye (50 µM) were added to the cells and incubated for 20 min at 37°C, 5 % CO_2_. Live cell imaging was performed by using Nikon Ti2 eclipse bright-field fluorescence microscope (40x magnification) including a CO_2_-O_2_ controller from Okolab. Measurements were started immediately after adding of stimuli. Time-lapse analysis was performed by taking pictures every 3 min for at least 2 hours. Image analysis was performed using NIS Elements from Nikon and Fiji analysis software.

### Luminex cytokine multiplex analysis

All samples were stored at −70 °C until testing. The samples were thawn at room temperature, vortexed, spun at 18,000 x g for 1 min to remove debris and the required sample volumes were removed for multiplex analysis according to the manufacturer’s recommendations. The samples were successively incubated with the capture microspheres, a multiplexed cocktail of biotinylated, reporter antibodies, and a streptavidin-phycoerythrin (PE) solution. Analysis was performed on a Luminex 100/200 instrument and the resulting data were interpreted using proprietary data analysis software (Myriad RBM). Analyte concentrations were determined using 4 and 5 parameter, weighted and non-weighted curve fitting algorithms included in the data analysis package.

### Cytometric bead array

A cytometric bead array was performed using the “Human inflammatory cytokine kit” from BD Bioscience (551811) and following the manufacturer’s instruction. 25 µl of samples and standards were added to 25 µl of the capturing bead mixture. Additionally, 25 µl of PE detection reagent was added to all tubes and incubated for 3 h at RT in the dark. Thereafter, 1 ml of wash buffer was added to each tube and centrifuged at 200 x g for 5 minutes. The supernatant was carefully removed and the pellet was resuspended in 300 µl wash buffer. Measurements were performed with the FACS Canto II from BD Bioscience operated using Diva software. Analysis was performed with Soft Flow FCAP Array v3 analysis software from BD Bioscience.

### Transwell experiments

PBMCs and neutrophils were always used from the same donor in each experiment. Transwell inserts were loaded with 100 µl of PBMC suspension (0.8 x 10^6^ cells/insert). In the lower chamber, either PMNs were seeded as described above (same plate size, same volume, same cell concentration) using polycarbonate, 24 well plates, 3 µm pores, Corning, 734-1570) and stimulated for 4 h with stimuli as indicated. Alternatively, media containing the stimuli only (i.e. no PMNs) or media containing only MIP-1β (30 and 150 pg /ml), IL-16 (300 and 1500 pg/ml) or SDF-1α (control, 100 ng/ml) and no PMNs were added. After 4 h, the lower compartment was harvested and FACS staining was performed as described above. The total number of migrated cells was acquired using counting beads (Biolegend, 424902) on a FACS Canto II (BD Bioscience) with Diva software. Analysis was performed using FlowJo V10 analysis software.

### Neutrophil elastase NETosis assay

Neutrophil extracellular trap formation was determined using the colorimetric NETosis Assay Kit from Cayman Chemicals based on the enzymatic activity of NET-associated neutrophil elastase. PMNs from various healthy donors were isolated as described above and stimulated with RNA-LL37 complex, or PMA and a calcium ionophore (A-23187) as positive controls for 1 or 3 hours. The assay was performed following the manufacturer’s instructions. The absorbance was then measured at 405 nm using a standard plate reader.

### Transient transfection of HEK293T cells

HEK293T cells were transiently transfected using the CaPO_4_ method as described (Colak et al., 2014). Cells were seeded in 24-well plates at a density of 14 x 10^5^ cells/ml 2-3 h prior to transfection. For the transfection of one well, 310 ng of plasmid DNA (100 ng TLR plasmid, 100 ng firefly luciferase NF-κB reporter, 10 ng *Renilla* luciferase control reporter, and 100 ng EGFP plasmid) was mixed with 1.2 µl of a 2 M CaCl_2_ solution and filled up with sterile endotoxin-free H_2_0 to obtain a total reaction volume of 10 µl. After the addition of 10 µl of 2X HBS solution (50 mM HEPES (pH 7.05), 10 mM KCl, 12 mM Glucose, 1.5 mM Na_2_HPO_4_), the mixture was then added to the cell suspension. As negative controls, TLR coding plasmids were replaced by empty vectors carrying the appropriate backbone of the TLR plasmids. After the addition of the transfection complexes, the cells were incubated either for 24 h followed by stimulation, or kept for 48 h without stimulation (MyD88 expression). For stimulation, the media was aspirated and replaced by fresh growth medium in which TNFα or the different TLR ligands (R848, CpG, RNA40) with or without IRS were diluted to appropriate concentrations. TLR8 activation with RNA40 was facilitated by complexation of the RNA with DOTAP (L787.1, Roth). RNA40 and DOTAP were first diluted in 1X HBS separately. Next, RNA40/HBS was diluted 1:3 in DOTAP/HBS. The solution was carefully mixed by pipetting up and down. After 15 min of incubation at RT, the mixture was 1:6.7 diluted in growth medium (with or without IRS) and finally dispensed (500 µl/well) into the wells containing transfected HEK293T cells. Each tested condition was measured in triplicates. The cells were stimulated and inhibited for 24 h at 37°C.

### Dual Luciferase Reporter Assay

After checking transfection efficiency via EGFP fluorescence microscopy, HEK293T supernatants were aspirated and 60 µl of 1X passive lysis buffer (E194A, Promega) added per well. The plate was then incubated for 15 min at RT on the plate shaker and subsequently stored at −80 °C for at least 15 minutes to facilitate complete cell lysis. After thawing, 60 µl were transferred into a V-bottom 96-well plate and centrifuged for 10 min at 2500 rpm and 4 °C to pellet cell debris. 10 µl supernatant were then transferred into a white microplate and each condition was measured in triplicates using the FLUOstar OPTIMA device (BMG Labtech). Firefly and Renilla luciferase activity were determined using the Promega Dual luciferase kit. Both enzyme activities were measured for 12.5 s with 24 intervals of 0.5 s, respectively. The data was analyzed by calculating the ratio of the two measured signals, thereby normalizing each firefly luciferase signal to its corresponding Renilla luciferase signal. The ratios were represented as the relative light units (RLU) of NF-κB activation.

### Statistics

Experimental data was analyzed using Excel 2010 (Microsoft) and/or GraphPad Prism 6 or 7, microscopy data with ImageJ/Fiji, flow cytometry data with FlowJo 10. In case of extreme values, outliers were statistically identified using the ROUT method at high (0.5%) stringency. Normal distribution in each group was always tested using the Shapiro-Wilk test first for the subsequent choice of a parametric (ANOVA, Student’s t-test) or non-parametric (e.g. Friedman, Mann-Whitney U or Wilcoxon) test. p-values (α=0.05) were then calculated and multiple testing was corrected for in Prism, as indicated in the figure legends. Values < 0.05 were generally considered statistically as significant and denoted by * throughout. Comparisons made to unstimulated control, unless indicated otherwise, denoted by brackets.

## Materials & Correspondence

Please address requests to: Alexander N. R. Weber, Interfaculty Institute for Cell Biology, Department of Immunology, University of Tübingen, Auf der Morgenstelle 15, 72076 Tübingen, Germany. Tel.: +49 7071 29 87623. Fax: +49 7071 29 4579.

## Supplementary Material

Is uploaded separately and includes 5 Supplementary figures and 6 supplementary tables.

## Authorship contributions

F.H., Z.B., S.D., D.E., T.K. and N.S.M. performed experiments; F.H., Z.B., S.D., D.E., N.S.M. and A.N.R.W. analyzed data; M.H., H. K., M.W.L., L.F., K.S., K.G. and T.E. were involved in sample and reagent acquisition; F.H., D.H., K.G. and T.E. contributed to the conceptual development of the study; F.H. and A.N.R.W. wrote the manuscript and all authors commented on an revised the manuscript; A.N.R.W. coordinated and supervised the entire study. None of the authors declare competing interests.

## Acknowledgements

We thank S. Pöschel, J. Berger, S. Haen, K. Preissner, O. Sorensen and A. Dalpke for provision of reagents, samples, technical support and/or helpful discussions and all healthy donors and patients for participation in our study. This study was supported by the Deutsche Forschungsgemeinschaft (German Research Foundation, DFG) CRC TR156 “The skin as an immune sensor and effector organ – Orchestrating local and systemic immunity”, the University of Tübingen and the University Hospital Tübingen.

## Abbreviations

APC: antigen-presenting cell
bRNA: bacterial ribonucleic acid
EGFP: Enhanced green fluorescence protein
HEK: Human embryonic kidney
HLA: Human leucocyte antigen
iODN: inhibitory oligonucleotides
MAMPs: Microbe-associated molecular pattern
MHC: Major histocompatibility complex
iODNs: Inhibitory oligodeoxynucleotides
IFN: Interferon
IL: Interleukin
KC: Keratinocytes
PASI: psoriasis area and severity
pDCs: Plasmacytoid dendritic cells
PMA: phorbol myristate acetate
PRR: Pattern recognition receptor
PMNs: Polymorphonuclear leukocytes
NE: Neutrophil elastase
NET: Neutrophil extracellular trap
SEM: Scanning electron microscopy
SLE: Systemic lupus erythematosus
TLR: Toll-like receptor
TNF: Tumor necrosis factor

---

**Figure S1.**
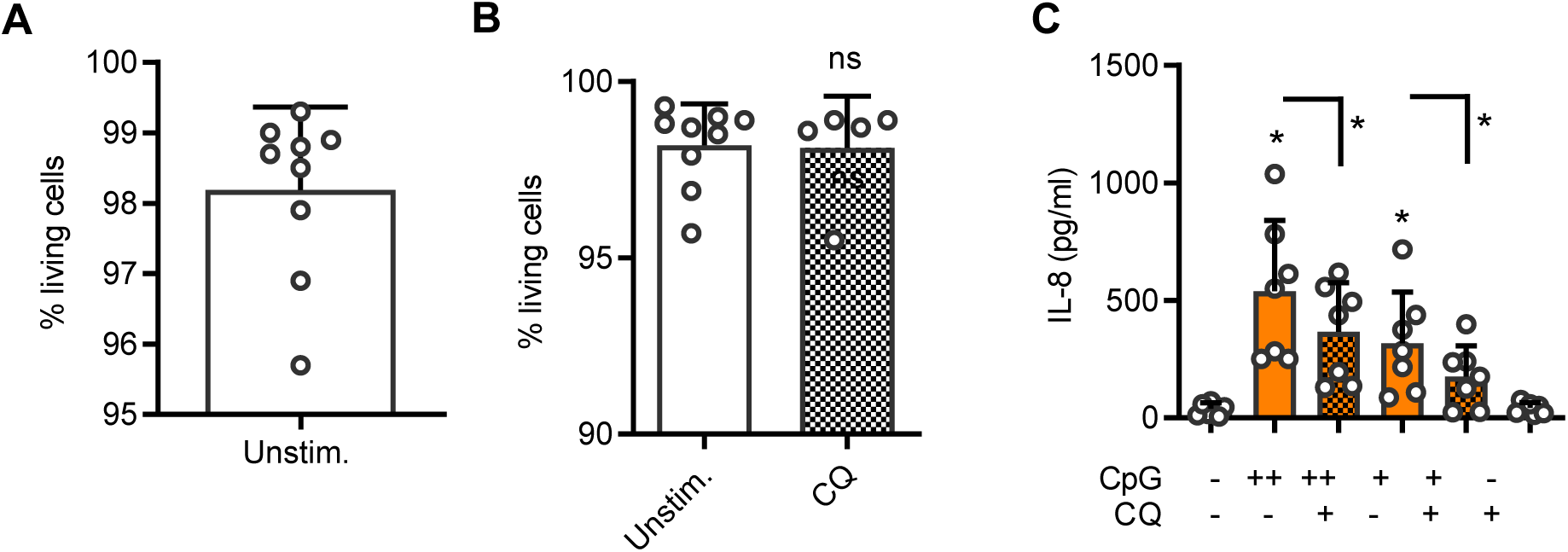

**Figure S2.**
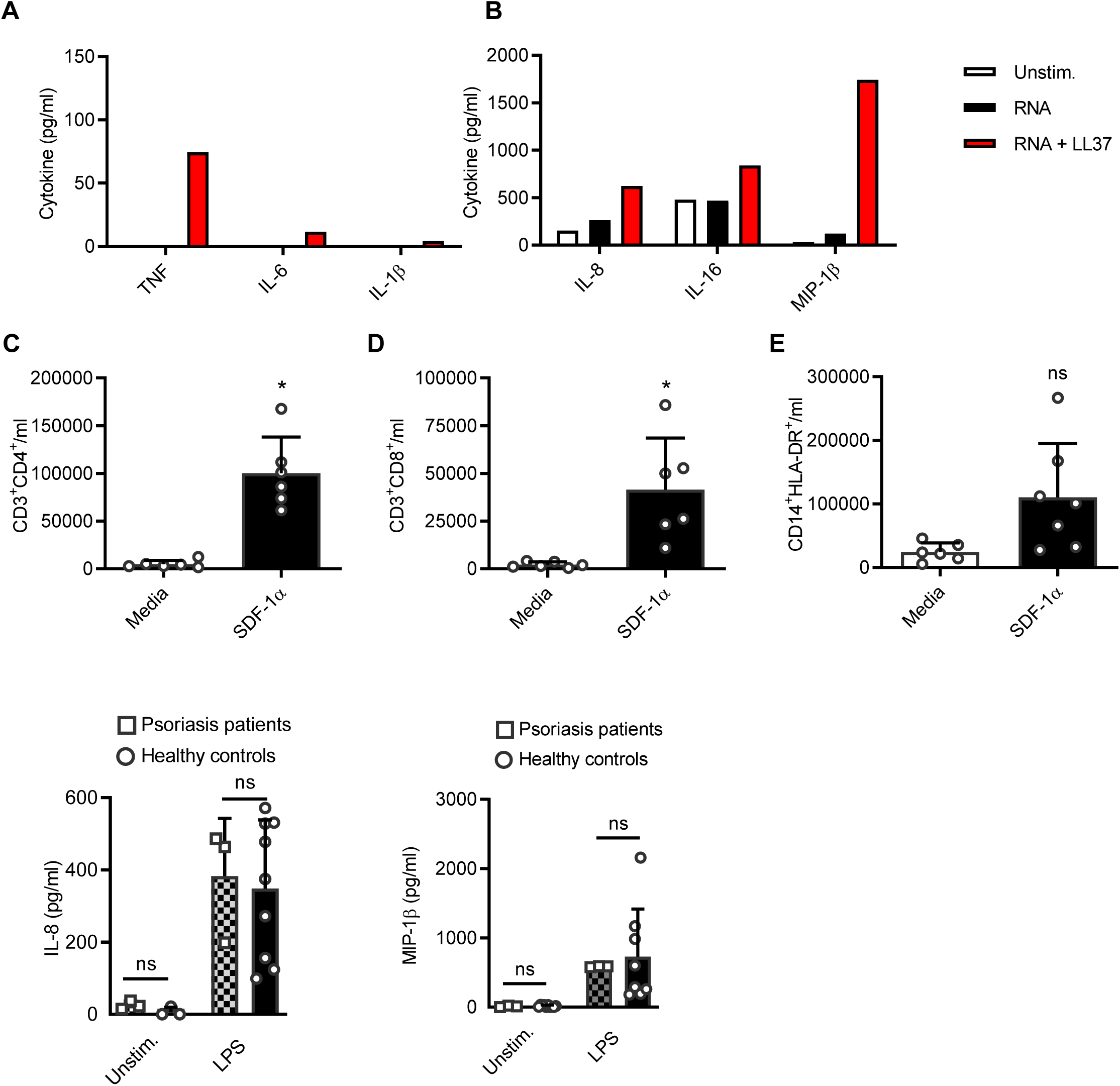

**Figure S3.**
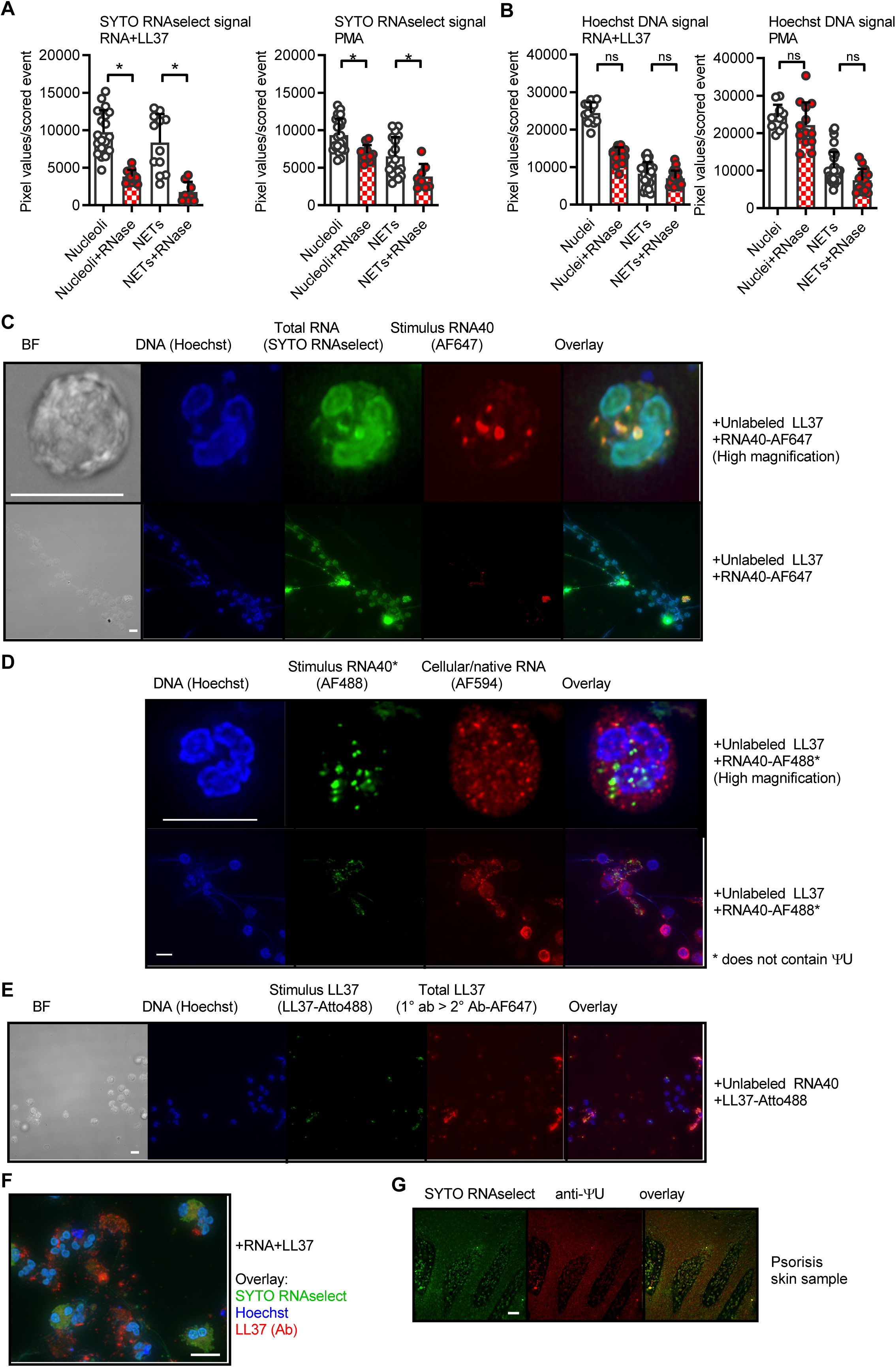

**Figure S4.**
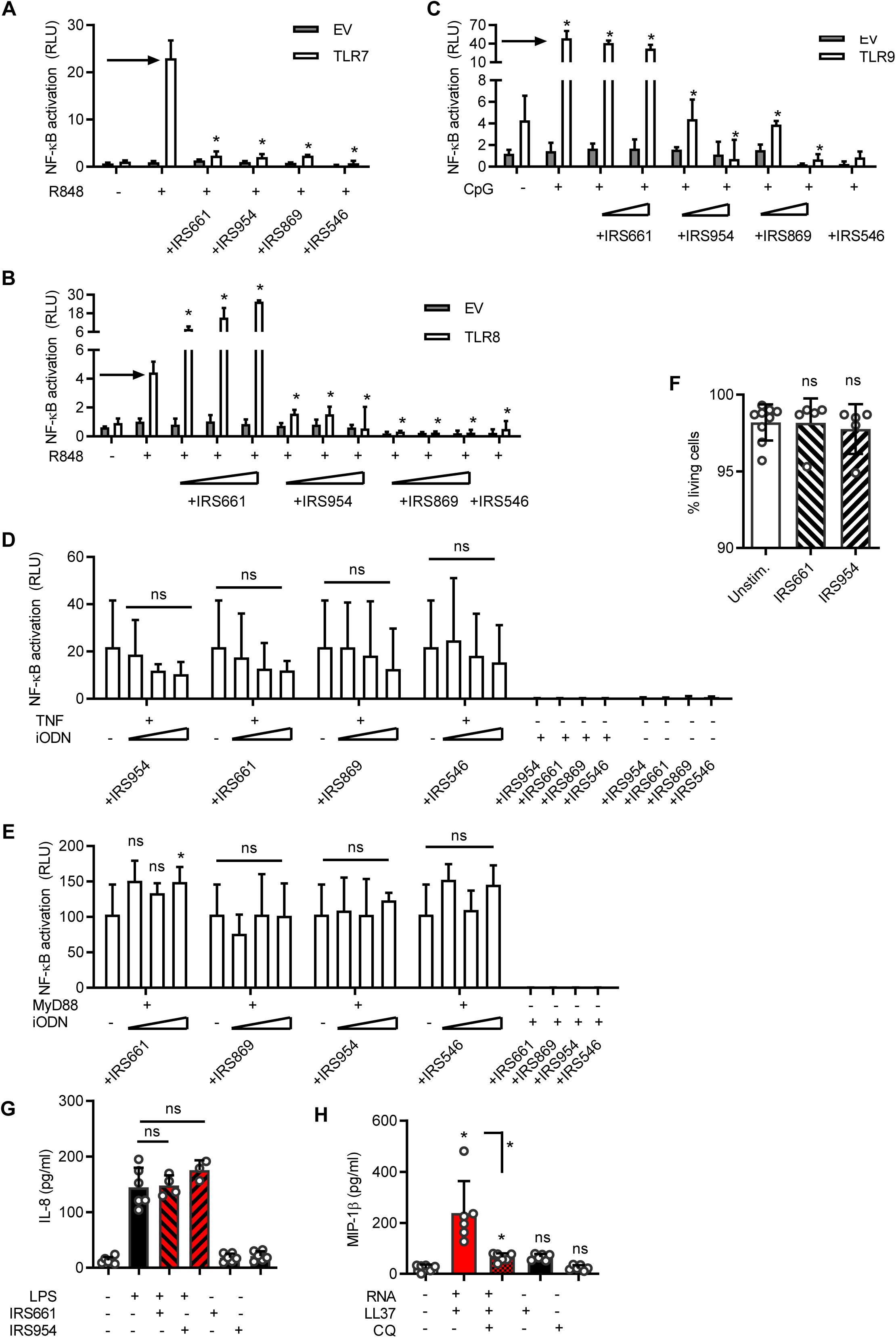

**Figure S5.**
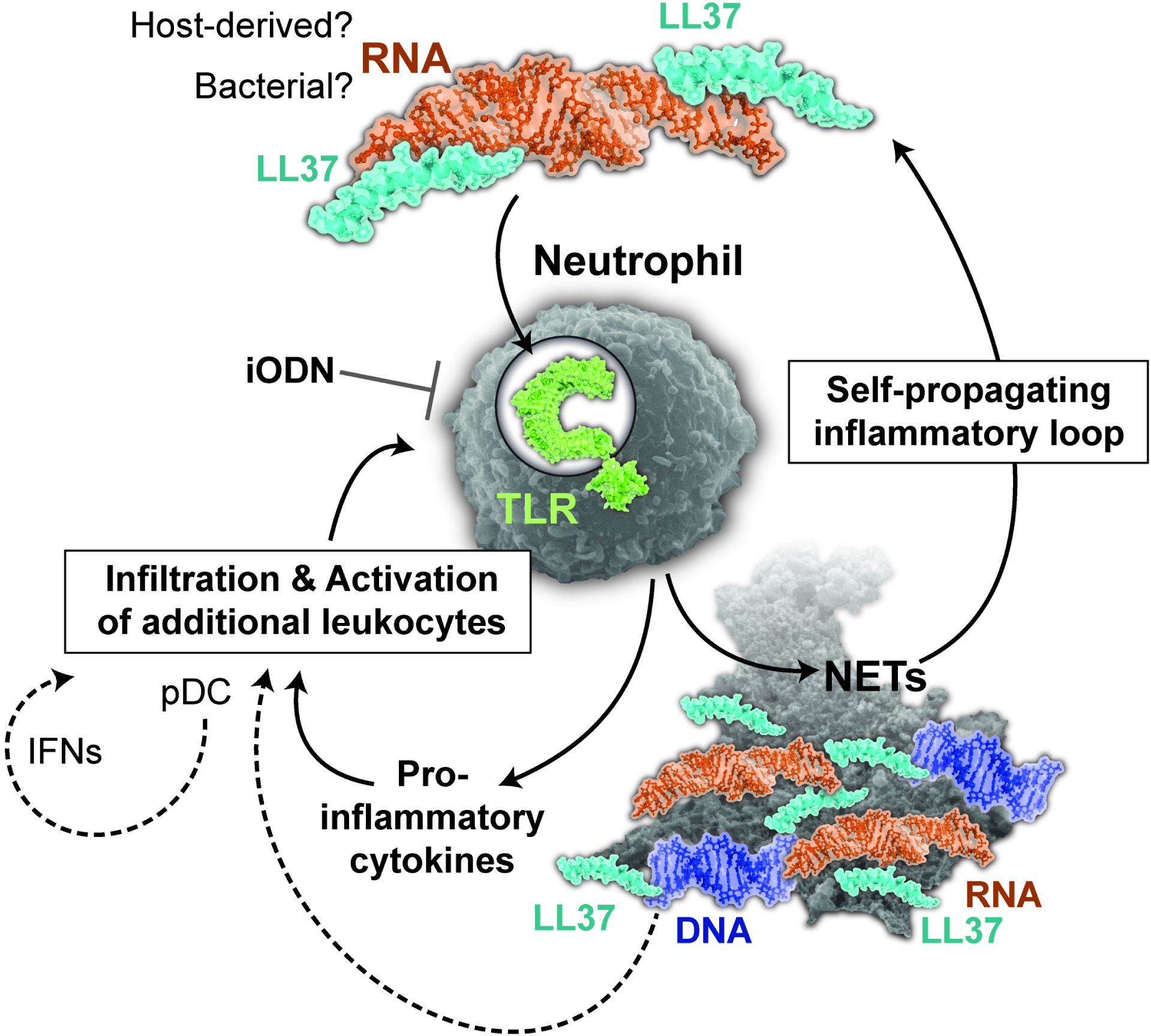

